# Evaluation of Ventilation at 10°C as the Optimal Storage Condition for Donor Lungs in a Murine Model

**DOI:** 10.1101/2025.08.11.669761

**Authors:** Morgan A. Hill, Megan Tennant, Bailey Watts, Carl Atkinson, Richard O’Neil, Kathryn E. Engelhardt, Barry C. Gibney

## Abstract

**Rationale:** Cold static preservation at 4°C is the clinical standard for donor lung storage but is limited to 6–8 hours of cold ischemia. Static storage at 10°C has been shown to extend ischemia times and improve lung health. Given that lungs can maintain aerobic metabolism ex vivo, we hypothesized that adding ventilation at 10°C would further prolong preservation by stimulating aerobic metabolism.

**Methods:** Lungs were procured from C57Bl/6 mice and then stored for 24h with ventilation at 10°C (n=4), statically at 10°C (n=4), or statically at 4°C (n=4). Respiratory mechanics were evaluated using a FlexiVent system. Cellular viability was assessed via flow cytometry. Complement shedding was evaluated by enzyme-linked immunosorbent assay. Histologic evidence of lung injury was assessed by H&E staining.

**Results:** Donor lungs stored with ventilation at 10°C exhibited significantly reduced histologic injury scores compared to static storage at 4°C (p = 0.0062). Ventilation also decreased complement C3 shedding (p < 0.01), apoptosis (p < 0.05), cytochrome c release (p = 0.0014), and ROS production (p = 0.0008) compared to statically stored lungs at 4°C and 10°C. Functionally, ventilated lungs demonstrated improved respiratory mechanics with lower airway resistance (p = 0.021) and increased compliance (p = 0.023) compared to static storage at 10°C.

**Conclusions:** Ventilating lungs at 10°C compared to static cold storage appears to result in healthier and more functional lung tissue and may extend the preservation times of donor organs for lung transplantation.

## INTRODUCTION

Recent changes to the lung donor allocation system^1^ have increased the number of lung transplants performed at the cost of increased travel distances for transplant centers^2^. Despite increased travel distance, lung recovery techniques have largely been unchanged, with cold static storage being the predominant method and the alternative being ex vivo lung perfusion (EVLP)^3^. Recent data suggests moderate hypothermia may extend cold ischemia time, attenuate donor lung injury, and improve cellular health within the lung allograft^4–6^. Regardless of lung storage, the technique for recovery is unchanged. Lung allografts are perfused with a cold flush – typically a low-potassium dextran solution^7^-while simultaneously ventilating the lungs with low-tidal volumes. This decreases atelectasis, which is associated with higher pulmonary vascular resistance and results in a heterogeneous distribution of perfusate. Following perfusion, the lung is inflated to 50% of lung capacity (or 15 cmH2O airway pressure) with 50% FiO2, and the trachea is clamped before placement in an ice cooler. While much focus has been placed on temperature and perfusion solutions, there has been less investigation into the role of stretch on the donor allograft.

During development, the lungs demonstrate significant sensitivity to stretch signals. Oligohydramnios, congenital diaphragmatic hernia, and phrenic nerve dysfunction^8–10^ – which all attenuate stretch signals – result in underdeveloped lungs. Excessive stretch signaling, such as with large tidal volume ventilation, exacerbates lung injury and leads to disordered alveolar growth^11–13^. Compensatory lung growth following pneumonectomy is well-described in many mammals^17^. This phenomenon can be attenuated by reducing cyclic stretch^18^ and appears to localize to subpleural regions of the lung – areas most subject to deformation^19,20^ – supporting the hypothesis that cyclic stretch is essential to alveologenesis. From a lung donation perspective, expanding the lung during recovery with oxygen allows for continued aerobic metabolism, preserved surfactant function, improved pulmonary compliance, and increased alveolar fluid clearance^14–16^. However, donor lungs are exposed to static stretch, and the role of cyclic stretch is unknown.

In this report, we applied cyclic stretch to a murine lung model to determine the effect of this stimulus on allograft health. After recovery, the lungs were subjected to static inflation or to continued room air ventilation at physiologic tidal volumes. We assessed mitochondrial and cellular health, histologic evidence of lung injury, and mechanical physiology in the context of each respective storage modality to assess if ventilation during storage results in more functional donor lungs.

## METHODS

### Animals and Surgical Procedure

This study was approved by the Committee of Animal Research following the National Institutes of Health Guide for Care and Use of Laboratory Animals. All personnel working with the animals had the required course training and certifications. C57Bl/6 mice were used for all experiments. The donor animal is induced with 5 parts per million (ppm) of isoflurane and maintained with 3 ppm of isoflurane via a nosecone. Depth of anesthesia is confirmed via toe pinch prior to the start of the procedure. The skin is divided with scissors from the xiphoid process to the jaw. The xyphoid process is retracted cephalad to expose the diaphragm. An incision is made on the right side of the diaphragm to collapse the lungs. The right and left ribs are then cut in the mid axillary line and retracted cephalad. 500 u/kg of heparin is then injected directly into the right atrium. The beating heart is then divided along the short axis to expose the right and left ventricular cavities. 50 ml/kg of Perfadex perfusate is delivered into the pulmonary artery from the right ventricle through the pulmonary valve using a gravity perfusion setup. After flushing is complete, the donor pneumonectomy is performed in standard fashion. Following donor pneumonectomy, the trachea is intubated with an 18-gauge Angiocath™ venous catheter (Beckton Dickenson, NJ, USA) and stored in one of three conditions: cold static storage at 4 °C (n=4), cold static storage at 10 °C (n=4), and ventilation storage at 10 °C (n=4). All lungs were stored in Perfadex solution for 24 hours.

### Cellular health

Murine lung tissue was harvested and enzymatically dissociated into a single cell suspension using the Lung Dissociation Kit (Miltenyi Biotec, North Rhine-Westphalia, Germany) according to the manufacturer’s instructions. Cellular viability was assessed with flow cytometry using Zombie UV fixable viability dye (ThermoFisher, MA, U.S.) to distinguish live and dead cells and Apotracker (BioLegend, CA, U.S.) to identify early apoptotic cells.

Mitochondrial health was evaluated via intracellular flow cytometry. Cells were stained with an anti-cytochrome c antibody (BioLegend, CA, U.S.) to assess mitochondrial membrane integrity and with MitoSOX Red (ThermoFisher, MA, U.S.) to detect mitochondrial superoxide production as an indicator of oxidative stress. All flow cytometry data were acquired on a CytoFLEX LX (Beckman Coulter, CA, U.S.) and analyzed using FlowJo software (BD Biosciences, NJ, U.S.). Gating strategies excluded doublets and debris based on forward and side scatter profiles.

### Histology

Lung tissue samples were embedded in paraffin after fixation in 10% buffered formalin for 48h, followed by 5μm sectioning and hematoxylin and eosin staining. The slides were then blindly reviewed and graded by two separate lung histopathologists using a previously described lung injury scale^22^. Briefly, lung injury was assessed based on four histologic criteria: white blood cell infiltration, fibrin exudates, alveolar hemorrhage, and capillary congestion. Each parameter was graded on a scale from 0 to 3, where 0 indicated absence, 1 mild, 2 moderate, and 3 severe involvement. Each animal’s cumulative injury score was calculated by summing the individual scores across all four parameters.

### Complement shedding

Murine C3 concentrations in the lung preservation solution were quantified via enzyme-linked immunosorbent assay (ELISA; Abcam, Cambridge, UK), performed in accordance with the manufacturer’s standardized protocol.

### Respiratory mechanics

*Ex vivo* pulmonary mechanics were evaluated using the FlexiVent small animal ventilator system (SCIREQ, Montreal, QC, Canada). Following 24 hours of storage, donor lungs were cannulated and connected to the FlexiVent platform for comprehensive respiratory function assessment. All assessments were completed at room temperature. Lung mechanics were quantified through a series of forced oscillation technique (FOT)-based perturbations. The snapshot perturbation maneuver was employed to derive key parameters, including airway resistance, dynamic compliance, tissue elastance, and hysterisivity. Pressure–volume relationships were assessed via ramp-style pressure-regulated perturbations to generate maximal pressure-volume loops.

## RESULTS

### Lung injury

Lung injury was quantified in a blinded manner using a validated histopathologic scoring system incorporating four criteria: leukocyte infiltration, fibrin deposition, alveolar hemorrhage, and capillary congestion. One-way ANOVA demonstrated a significant effect of the storage condition on cumulative lung injury scores (p= 0.0079) (**Figure 1**). Tukey analysis revealed that lungs ventilated at 10°C exhibited significantly reduced histologic injury compared to those stored statically at 4°C (p = 0.0062). Although ventilated lungs also demonstrated lower injury scores relative to static storage at 10°C, this difference did not reach statistical significance (p = 0.4238).

**Figure 1:**
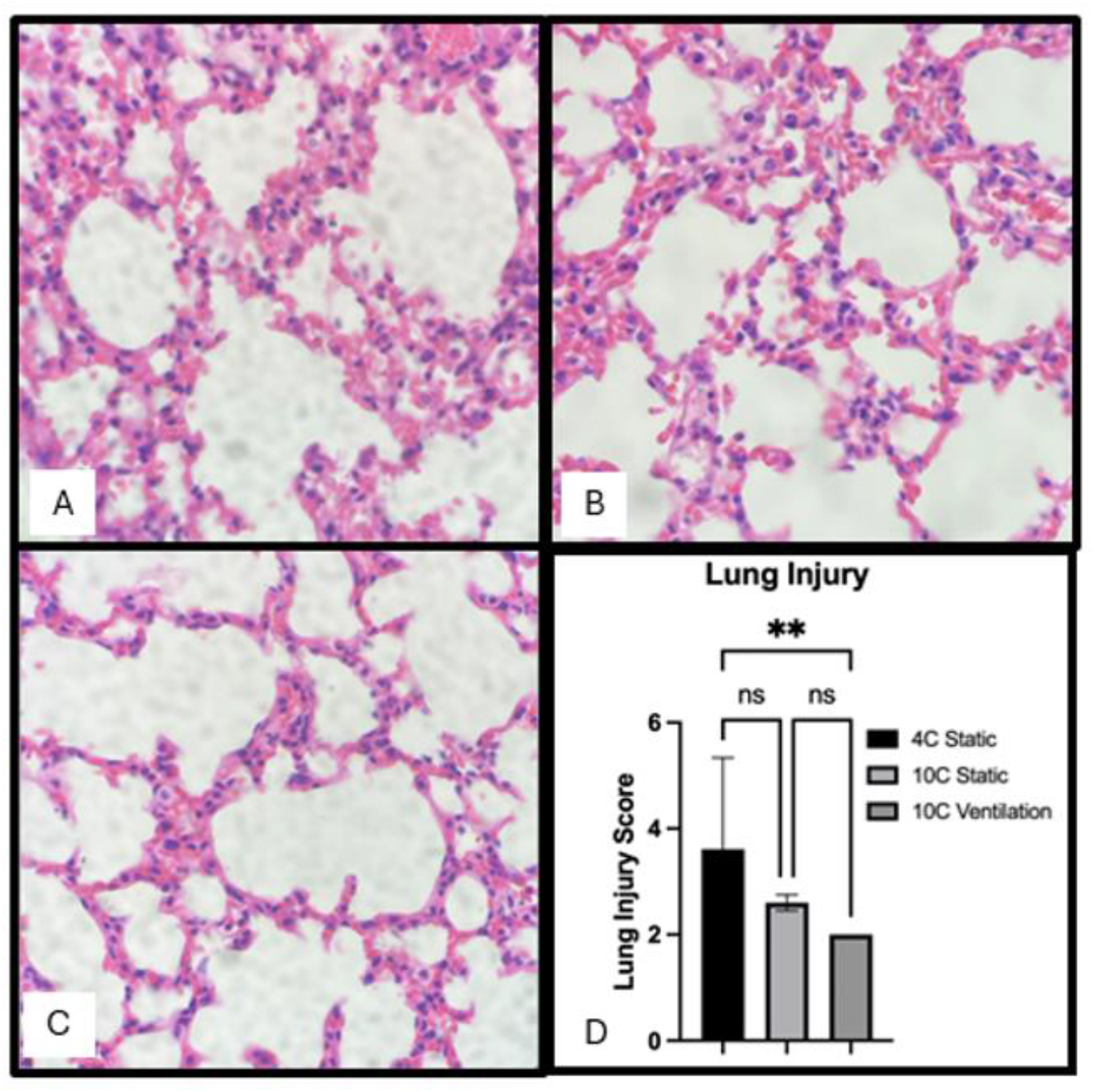
Recovered murine lung allografts were assessed for: leukocyte infiltration, fibrin deposition, alveolar hemorrhage and capillary congestion to determine the lung injury score. Lungs stored with the addition of ventilation demonstrated significantly less lung injury. Representative hematoxylin and eosin-stained lung images for (A) lung allografts stored for 24 hours at 4^°^C static inflation, (B) 10^°^C static inflation, (C) 10^°^C ventilated with room air at tidal volume ventilation, and (D) Quantification of Lung Injury Score in each group.. One-way ANOVA demonstrated a significant effect of the storage condition on cumulative lung injury scores (p= 0.0079). Lungs ventilated at 10°C exhibited significantly reduced histologic injury compared to those stored statically at 4°C (p = 0.0062).

### Complement shedding

Complement C3 concentrations in the lung preservation solution were quantified via ELISA. Ventilated donor lungs stored at 10°C exhibited significantly reduced C3 shedding (84.3 ± 33.8 ng/mL) compared to lungs stored statically at 4°C (390.3 ± 129.5 ng/mL; p = 0.0102) and 10°C (517.0 ± 13.4 ng/mL; p = 0.0011) **(Figure 2)**.

**Figure 2:**
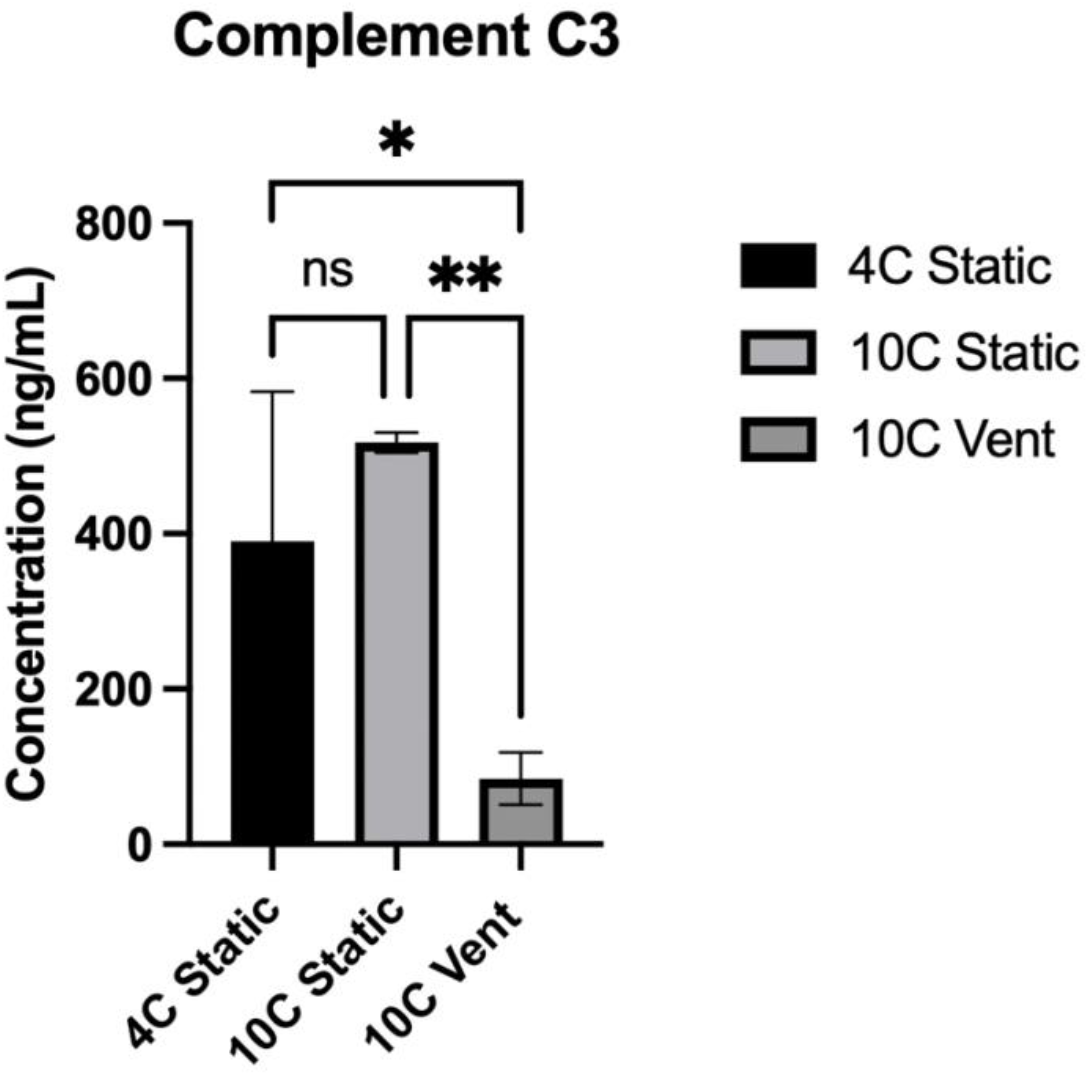
Lung allografts that were stored with tidal volume ventilation demonstrated significantly lower levels of Complement C3 in the storage perfusate when compared to lungs stored at static inflation at 4°C (p=0.0102) and 10°C (p=0.0011).

### Cellular health

Donor lungs were enzymatically dissociated and analyzed by flow cytometry to assess cellular viability and apoptosis, utilizing both live/dead discrimination and apoptotic staining. Ventilated lungs stored at 10°C demonstrated a significantly lower proportion of apoptotic cells (45.2% ± 2.25%) compared to static storage at 4°C (55.7% ± 3.61%; p = 0.0016) and 10°C (51.2% ± 2.65%; p = 0.0386). Although a higher percentage of viable cells was observed in the ventilated group, this difference did not reach statistical significance (p = 0.18) **(Figure 3)**. Mitochondrial integrity was assessed by quantifying cytochrome c release—an indicator of mitochondrial outer membrane permeabilization during apoptosis—and intracellular reactive oxygen species (ROS) generation. Donor lungs ventilated at 10°C exhibited significantly reduced cytochrome c levels (20.0 ± 6.17 MFI) compared to lungs stored statically at both 4°C (50.35 ± 8.77 MFI) and 10°C (37.25 ± 8.49 MFI; p = 0.0014). Storage condition also significantly influenced ROS production across groups (P = 0.0011). Tukey analysis revealed that ventilated lungs at 10°C generated significantly less ROS (1819 ± 231.1 MFI) than those stored statically at 4°C (3121 ± 360.9 MFI; p = 0.0008), with a non-significant trend toward reduced ROS relative to static 10°C storage (2440 ± 362.9 MFI; p = 0.057) **(Figure 4)**.

**Figure 3:**
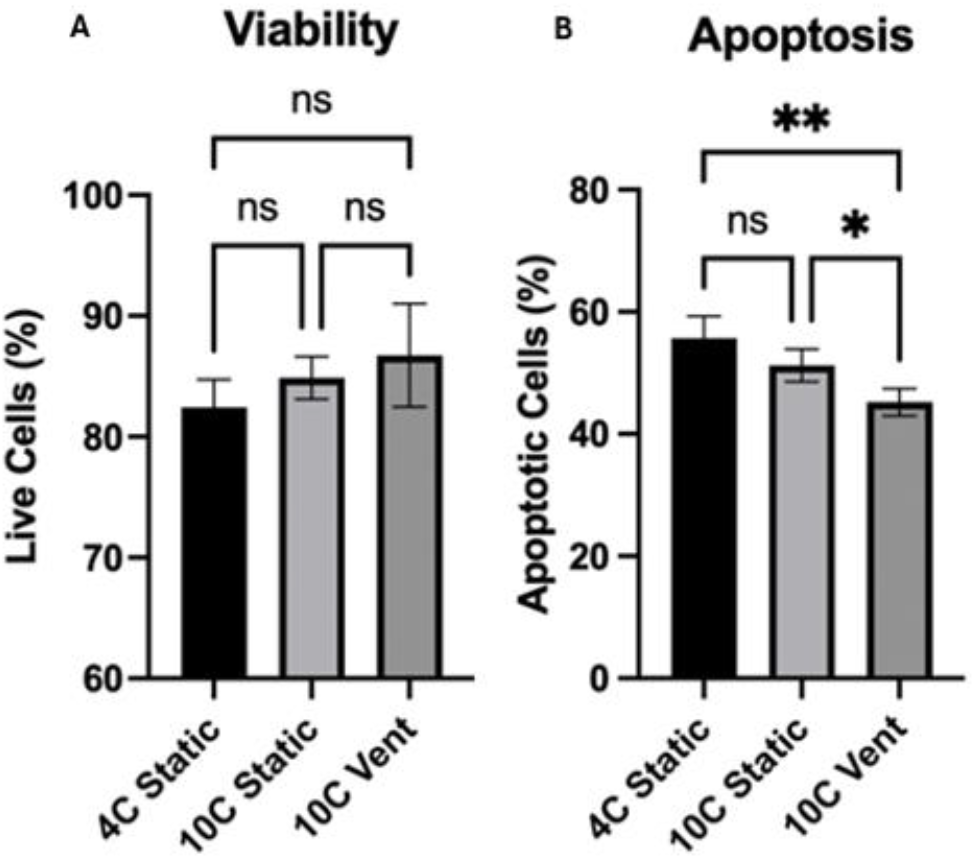
Lungs were digested after 24 hours of storage and assessed with live/dead staining (Zombie UV) and apoptosis (Apotracker). (A) A non-significant trend in improved cellular viability was seen with the addition of ventilation to the stored lungs. (B) Ventilation significantly decreased the percentage of apoptotic cells compared to static storage at 4°C (p = 0.0016) and 10°C (p = 0.0386).

**Figure 4:**
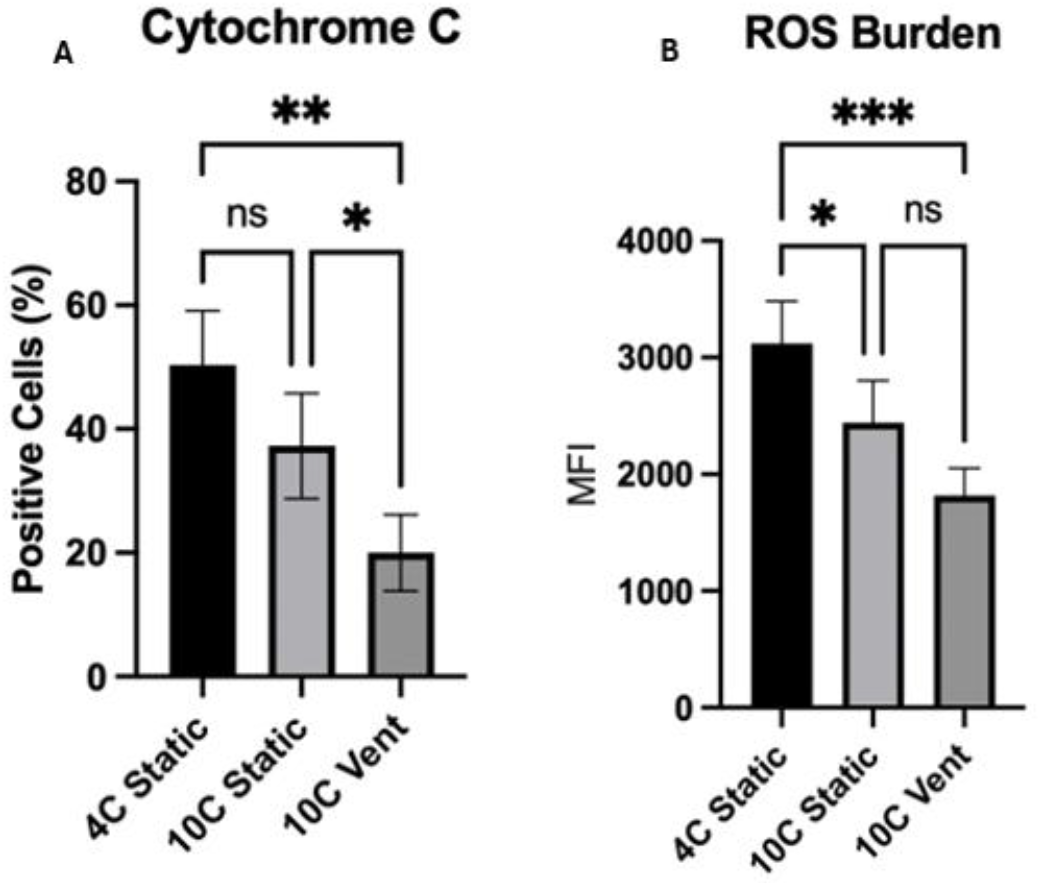
Mitochondrial health was assessed using intracellular flow cytometry evaluation staining for cytochrome C and superoxide production. (A) The addition of ventilation significantly reduced cytochrome C production (p=0.0014). (B) Temperature significantly reduced superoxide production (p=0.0011), though no significant difference was seen between static storage and ventilation at 10^°^C (p=0.057).

### Respiratory mechanics

Pulmonary function was evaluated using the FlexiVent small animal ventilator system to characterize the impact of preservation strategy on respiratory mechanics. Ventilated lungs stored at 10°C demonstrated significantly reduced airway resistance (0.88 ± 0.46 cmH_2_O·s/mL) compared to statically stored lungs at 10°C (3.06 ± 0.84 cmH_2_O·s/mL; P = 0.021), along with a significant increase in dynamic compliance (0.016 ± 0.003 mL/cmH_2_O vs. 0.006 ± 0.0008 mL/cmH_2_O; P = 0.023) **(Figure 5)**. Although differences in peripheral lung mechanics— specifically tissue elastance, damping, and hysterisivity—did not reach statistical significance, ventilated lungs exhibited a consistent trend toward improved values across these parameters **(Figure 5)**.

**Figure 5:**
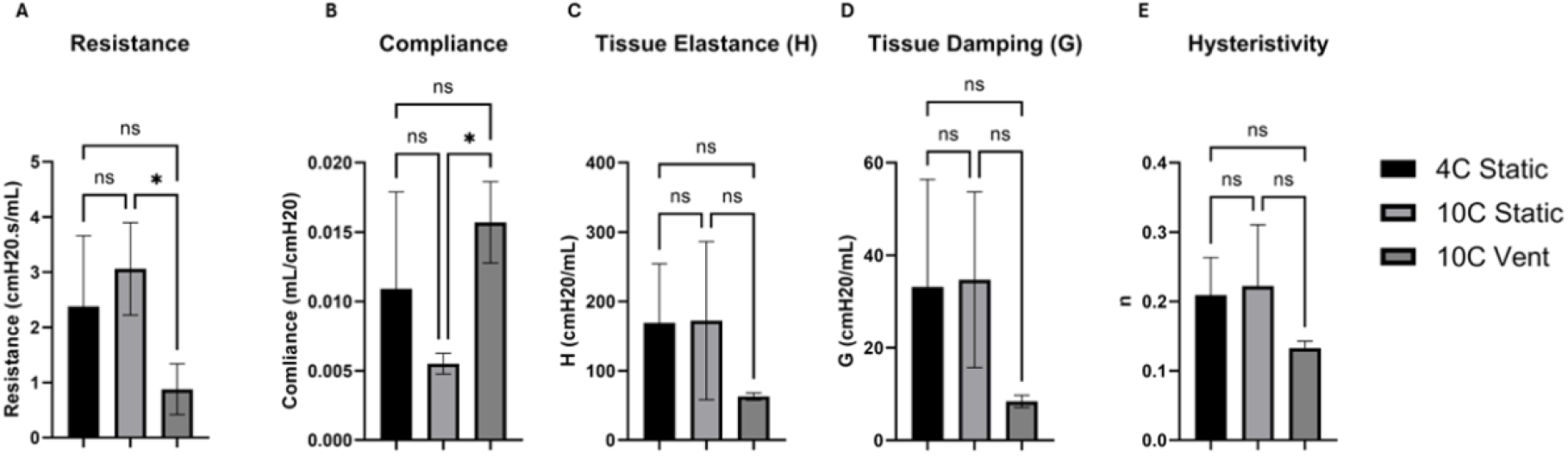
Lung mechanics were assessed after 24 hours storage using the FlexiVent small animal ventilator to evaluate single-compartment mechanics and measures from the forced oscillation maneuver. (A) Single-compartment airway resistance in ventilated lungs was significantly decreased (p=0.021) and (B) pulmonary compliance significantly improved (p=0.023) compared to statically stored lungs at 10°C. When evaluating the lung using forced oscillation, (C) Elastance and (D) Damping were reduced in the lungs subjected to ventilation, though not significantly (p=0.157, p=0.106). (E) Likewise, hysteresivity was reduced but not significantly (p=0.132).

## DISCUSSION

In this report, the addition of normal tidal volume ventilation to recovered murine lungs produced five principal findings: 1) Cell viability increased, and apoptosis decreased. 2) Mitochondrial health significantly improved, with ventilated cells demonstrating lower levels of cytochrome C and reduced reactive oxygen species. 3) Ventilated allografts exhibited significantly less lung injury when assessed with H&E staining. 4) The storage perfusate showed a significant decrease in complement shedding. 5) Pulmonary function improved in donor lungs stored with ventilation. Our data demonstrate that donor lungs benefit from ventilation during cold storage.

Alveolar recruitment has long been demonstrated as advantageous following lung recovery. In an experiment assessing the effect of alveolar recruitment on ischemia-reperfusion, DeCampos et al. compared the effects of inflation to TLC with those of prolonged tidal volume ventilation against standard reperfusion in a rat model. The group showed significant improvement in pO2, decreased shunt fraction, and reduced peak airway pressure. Pulmonary edema was also significantly improved with alveolar recruitment. Importantly, any alveolar recruitment was beneficial, as no difference was seen between TLC inflation and 10 minutes of ventilation^21^. Consistent with these findings, our data show that application of ventilation during lung preservation at 10°C led to improved respiratory mechanics, specifically demonstrating significantly lower airway resistance and increased dynamic compliance compared to lungs stored statically. These results suggest that application of non-injurious cyclic stretch during storage may confer functional benefits to the donor lung, likely through sustained alveolar recruitment and mitigation of atelectasis-related injury.

We also observed that cyclic stretch applied via ventilation during storage improved mitochondrial health, a finding that is likely attributable to enhanced mitochondrial biogenesis. In support of this mechanism, Kim et al. demonstrated that cyclic stretch upregulates key regulators of mitochondrial biogenesis and oxidative phosphorylation—such as PGC-1α, TFAM, and ERRα—leading to increased mitochondrial mass and ATP production in cardiac myocytes^23^. In the context of pulmonary epithelial cells, McAdams and colleagues reported that non-injurious cyclic stretch under hyperoxic conditions reduced superoxide accumulation and preserved cell viability, suggesting that mechanical stretch may suppress ROS production directly or upregulate endogenous antioxidant defenses^24^. Similarly, Zhou et al. showed that controlled lung inflation during preservation elevated superoxide dismutase (SOD) activity and reduced oxidative stress markers, further supporting the role of mechanical forces in redox homeostasis^25^. Collectively, these findings reinforce a mechanistic paradigm in which cyclic stretch during lung preservation enhances mitochondrial biogenesis and function, thereby attenuating oxidative injury through improved mitochondrial quality control and redox regulation.

We also found that ventilated lungs stored at 10°C shed significantly less complement C3 compared to statically stored lungs, suggesting a potential reduction in complement activation under this preservation strategy. Complement activation has emerged as a key contributor to primary graft dysfunction following lung transplantation^26-28^. Prior studies have demonstrated that complement split products, such as C3d and C4d, deposit in the pulmonary microvasculature early after transplantation, particularly in cases complicated by PGD. Specifically, Westall et al. identified widespread septal capillary deposition of C3d and C4d in lung allografts within the first three months post-transplant, correlating with early graft injury^27^. More recently, Kulkarni et al. showed that levels of various complement activation fragments, including sC4d, sC5b-9, C1q, C2, C4, and C4b, were significantly elevated in bronchoalveolar lavage fluid from patients with severe PGD, implicating activation of all three complement pathways^28^. Furthermore, inhibition of C3 activation in a murine transplant model has been shown to protect against ischemia-reperfusion injury and lung injury, underscoring the pathogenic role of complement in early graft dysfunction^29^. In light of these findings, our study demonstrated that donor lungs ventilated at 10°C during preservation shed significantly less C3 compared to statically stored lungs, suggesting that ventilation at sub-normothermic temperatures may mitigate complement activation during storage and potentially reduce early graft injury.

This study presents a novel method of lung preservation and demonstrates consistent improvements in allograft health based on analysis of several key physiological parameters. However, these results should be considered in the context of certain limitations relevant to the models used. Specifically, while the murine model provides a controlled platform for mechanistic investigation, it does not fully recapitulate the anatomic and immunologic complexity of human lungs. Additionally, our study focused on pre-transplant allograft quality without assessing post-transplant function, leaving the long-term impact of ventilated storage on graft performance unresolved. Follow-up studies will be designed to leverage additional conditions beyond the use of physiologic tidal volumes at 10°C, which will further clarify the optimal parameters for stretch and ventilation during allograft storage. Specifically, these future studies will explore the optimal combination of ventilation parameters—such as tidal volume, rate, pressure, and oxygen concentration—across different preservation temperatures. In addition, further investigating the cellular and molecular pathways influenced by cyclic stretch, and expanding this work to include transcriptomic or proteomic profiling could further clarify the mechanisms by which ventilation preserves graft quality. Finally, future studies using large animal and transplant models will be critical to determine the clinical translatability of these findings. Together, these directions aim to refine and validate a ventilation-based preservation strategy that could meaningfully enhance donor lung utilization and post-transplant outcomes.

## ACKNOWLEDGEMENTS

We would like to acknowledge Konrad Rajab, MD, for his contributions during the early stages of this project, including concept development and support in obtaining initial resources.

